# Three-dimensional label-free observation of individual bacteria upon antibiotic treatment using optical diffraction tomography

**DOI:** 10.1101/775346

**Authors:** Jeonghun Oh, Jea Sung Ryu, Moosung Lee, Jaehwang Jung, Seung yun Han, Hyun Jung Chung, Yongkeun Park

## Abstract

Measuring alterations in bacteria upon antibiotic application is important for basic studies in microbiology, drug discovery, and clinical diagnosis, and disease treatment. However, imaging and 3D time-lapse response analysis of individual bacteria upon antibiotic application remain largely unexplored mainly due to limitations in imaging techniques. Here, we present a method to systematically investigate the alterations in individual bacteria in 3D and quantitatively analyze the effects of antibiotics. Using optical diffraction tomography, *in-situ* responses of *Escherichia coli* and *Bacillus subtilis* to various concentrations of ampicillin were investigated in a label-free and quantitative manner. The presented method reconstructs the dynamic changes in the 3D refractive-index distributions of living bacteria in response to antibiotics at sub-micrometer spatial resolution.

## 1. Introduction

Understanding how antibiotics induce bacterial cell death is essential in the study of microbiology and also to general healthcare in order to help develop, improve, and apply treatment to patients [1]. Over the past years, the mechanism of antibiotics has been studied intensively along with the use of new multiple classes of antibiotics that have succeeded in treating infectious diseases [2]. However, the widespread use of antibiotics has resulted in emergence of drug-resistant bacteria, evoking a significant challenge to the use of antibiotics [3, 4]. Since the mainstream approach to overcome this crisis primarily relies on the discovery and development of newer, more efficient antibiotics, understanding how bacteria respond to antibiotics has become more critical.

A basic approach to understanding the mechanism of antibiotics is to observe the phenotypic or physical response of bacteria to antibiotics. Phenotypic imaging has been one of the most popular diagnostic methods to achieve rapid antimicrobial susceptibility testing [5–10], as it provides an easy, low cost, real-time single-cell level examination. Furthermore, phenotypic analysis of bacteria has been exploited to extract useful information in various fields such as epidemiology and systems biology [11]. Although the phenotypic response of bacteria to β-lactam antibiotics has been studied [12, 13], detailed phenotypic aspects have been less explored with little available research on the changes in physical quantities such as cell mass, density, and volume. These parameters are direct indicators of cell survival and growth [14, 15], which can provide fundamental insights into the physical and chemical mechanism of bacterial response to antibiotics.

To revisit these unattended but physiologically relevant quantities of bacteria, we need an imaging modality that quantifies the parameters mentioned above. However, conventional two-dimensional (2D) microscopic techniques, such as phase contrast microscopy or differential interference contrast (DIC) microscopy, are not suitable for this purpose, because they are unable to measure three-dimensional (3D) properties. Fluorescence microscopy, such as scanning confocal fluorescence microscopy, allows 3D molecular imaging of cells [16], but the use of fluorescent dyes or proteins have limitations for studying the dynamic alterations in bacteria over a long time due to issues of photo-bleaching and photo-toxicity [17]. Scanning electron microscopy or atomic force microscopy is also not suitable for assessing the dynamics of live bacteria with high throughput [18, 19].

To investigate the aforementioned parameters, optical diffraction tomography (ODT), one of the 3D quantitative phase imaging (QPI) techniques, has several potentials [20]. ODT provides a non-invasive and non-contact 3D optical imaging [21]. ODT reconstructs the 3D refractive index (RI) maps of optically transparent samples based on interferometric imaging at multiple illumination angles. Because RI distributions serve as both intrinsic cell markers and indicators of protein densities, ODT meets the demands of quantitative phenotypic imaging of unlabeled live cells. Thus, its applications have been rapidly expanded for investigations on hematology [22], microalgae [23], immunology [24], infectious diseases [25, 26], plant biology [27], cell biology [27, 28], neuroscience [29], and drug discovery [30]. Some bacterial studies have also used ODT [31], but its application for bacterial microbiology is in a nascent stage.

In this study, we investigated the 3D bacterial response to antibiotics by exploiting ODT. Our method provides a label-free, real-time analysis of the phenotypic changes in bacterial cells at given antibiotic conditions. The quantitative analysis of 3D RI maps enables examination of the physical changes in cells induced by antibiotic agents. We demonstrate the capability of ODT for imaging the antibiotic response of bacteria in 3D by investigating the response of *Escherichia coli* and *Bacillus subtilis* to a β-lactam antibiotic agent, ampicillin. The dynamics of bacteria were observed by mapping the 3D RI distribution in real time, and the changes in morphological and biochemical parameters were analyzed. With its non-invasive, label-free, and quantitative analysis, our method provides a useful tool for studying and developing antibiotic agents against clinical pathogens that threaten public health.

## 2. Methods

### 2.1 Optical diffraction tomography

To measure the 3D RI tomogram of individual bacteria, we employed optical diffraction tomography based on a Mach-Zehnder interferometer equipped with a digital micromirror device (DMD) [Fig. 1(a)].

**Fig. 1.**
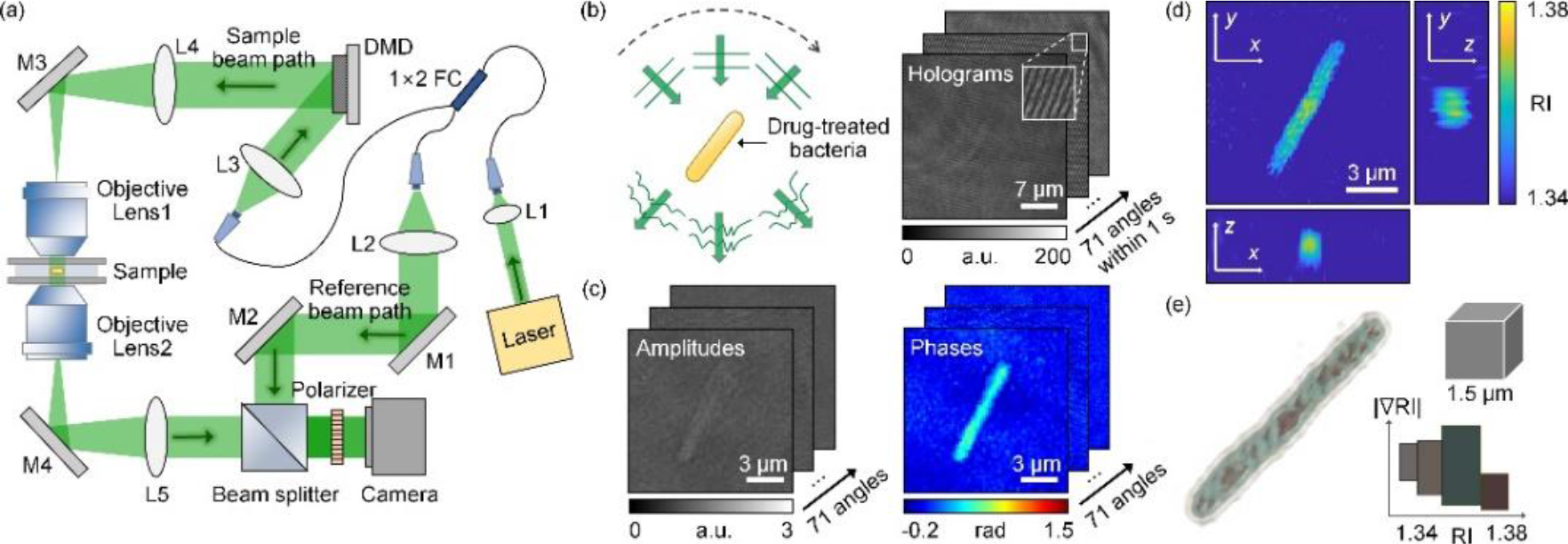
Measurements of the 3D RI distribution. (a) Experimental setup. M1–M4, mirror; L1–L5, lens; FC, fiber coupler; DMD, digital micromirror device. (b) Sequential illumination scanning for bacteria at 71 illumination angles and the corresponding 71 holograms. (c) Images of the retrieved amplitudes and phases corresponding to the holograms. (d) Three-section view of the reconstructed RI distribution. (e) 3D rendered image of bacteria. The color map shows various colors depending on the value of RI and its gradient value.

A coherent laser (*λ* = 532 nm in a vacuum; Samba™, Cobolt Inc., Sweden) beam was split into a reference beam and sample beam using a 1 × 2 fiber coupler. The sample beam passed through 4-*f* relay lenses (L1–3), and was diffracted by a DMD (DLPLCR6500EVM, Texas Instruments Inc., USA). A DMD controlled the angle of the sample beam impinging onto a sample, by displaying digital grating patterns [32]. The time-multiplexing illumination technique was utilized to eliminate the unwanted diffractions of a DMD [33]. The first-order diffraction beam illuminated a sample with a specific incident angle controlled by the pattern projected onto the DMD. The beam diffracted from the DMD passes through a lens (L4) and a condenser lens (NA = 1.2, water immersion, UPLSAPO 60XW, Olympus Inc.) in order to magnify the incident angle. The scattered light from the sample was collected by an objective lens (NA = 1.2, water immersion, UPLSAPO 60XW, Olympus Inc.), and projected to the image plane. At the image plane, the sample beam and reference beam interfere and generate a spatially modulated hologram pattern, which is recorded using a camera (FL3-U3–13Y3M-C, Point Grey Research Inc.). To combine two beams and generate the interference pattern with the maximum contrast, a beam splitter and a linear polarizer were used, respectively. The beam impinging onto the sample was controlled to have the maximum elevation angle limited by the NA of the condenser lens and was scanned along a full azimuthal angle with 71 steps [Fig. 1(b)]. The total acquisition time for 71 holograms was approximately 1 s (15 ms per hologram).

### 2.2 3D RI tomogram reconstruction

The 3D RI distribution of a sample is reconstructed from the measured multiple 2D holograms using the principle of optical diffraction tomography [34, 35]. For each hologram pattern, amplitude and phase delay maps were retrieved using the field retrieval algorithm [Fig. 1(c)] [36, 37]. Then, from these retrieved optical field maps with various angles, the 3D RI distribution was determined [Fig. 1(d)]. ODT solves an inverse scattering problem for monochromatic light propagating in a linear, isotropic, and nonmagnetic medium, which satisfies the 3D inhomogeneous Helmholtz equation [34]. For a slowly varying scattered complex phase, the Rytov approximation was applied, which has shown better reconstruction performances than the Born approximation [38, 39]. According to the Fourier diffraction theorem, each optical field is mapped onto a sphere in the 3D Fourier space [40]. Then, the inverse Fourier transform of the 3D Fourier space provides the RI distribution of the sample [41]. The theoretically calculated lateral and axial resolution of the used optical imaging system is 124 nm and 397 nm, respectively [32, 42]. Due to the limited NAs of both the condenser and the objective lens, side scattering signals are not collected, which is known as the missing cone problem. To resolve this missing cone problem, the regularization algorithm based on non-negativity criteria was used [43–45].

### 2.3 Analysis of cell parameters and visualization methods

To measure cell volumes, the voxels were selected which have an RI higher than a specific value determined by masking the sample region at the initial time for each cell and averaging the RI values therein. Because the RI value of cell cytoplasm is linearly related to its protein concentration, local protein concentration of a cell can be retrieved from the measured 3D RI tomogram [46, 47]. The concentration at each point of the bacteria was determined from RI increment (RII), an increment of an RI in a solution per an increment of concentration in a solute. We assumed that the value of RII is constant inside bacteria, *i.e. n*(**r**) − *n*_medium_ = (RII) × *C*(**r**), where *n* is an RI inside the bacteria, *n*_medium_ is the RI of surrounding medium, and *C* is the protein concentration. Therefore, the concentration distribution inside bacteria can be acquired using an RI distribution. In addition, the RII of 0.185 mL/g is known as a typical value for proteins [42, 46]. Because many bacteria have proteinaceous organelles [48], it is reasonable that the typical value of RII for proteins could be used in our analysis. The temporal changes in the cellular dry mass are calculated as the total sum of the concentration at each voxel multiplied by its volume. Cytoplasm concentration was calculated by averaging the concentrations at the internal voxels of bacteria.

The 3D rendered image was obtained using commercial software (TomoStudio™, Tomocube Inc., Republic of Korea), which delineates the overall RI distributions of bacteria more clearly [Fig. 1(e)]. The different colors indicate the presence of subcellular organelles in bacteria with different RI values, suggesting that our technique provides sub-micrometer spatial resolution to investigate individual bacterial cells.

### 2.4 Sample preparation

Bacterial strains *Escherichia coli (E. coli*; #1682) and *Bacillus subtilis* (*B. subtilis*; #1021) were purchased from Korean Collection for Type Cultures. *E. coli* cells were cultured in Tryptic soy broth (TSB) and *B. subtilis* were cultured in Nutrient broth (NB). Both bacteria were cultured to the concentration of 1.0−5.0 × 10^8^ cells/mL to obtain activated bacteria using a nanophotometer (Implen). Centrifugation to wash the cells was repeated twice under the general condition of 8,000 × *g* for 5 minutes at 4°C in 200 μL of PBS solution. Cells were then resuspended in TSB and NB to conduct imaging for the bacteria. Ten microliters of the resuspended solution were placed between two coverslips and then measured using the optical imaging system. All the measurements were performed at room temperature (25°C).

## 3. Results

In order to systematically investigate the alterations in individual bacteria upon treatment with antibiotics, we measured the 3D RI tomograms of *E. coli* and *B. subtilis* using ODT, and analyzed the measured images.

### 3.1 Changes in a bacterial cell revealed by time-lapse 3D imaging

We first validated the 3D dynamic imaging capability of ODT by visualizing bacterial growth in the absence of antibiotics. We imaged the dynamics of *E. coli* in growth medium with a time interval of 30 minutes up to 6 hours. Both the cross-sections and rendered images of the measured 3D RI tomograms clarify the growth and division of individual *E. coli* [Figs. 2(a) and 2(b)]. Prominent cell division was observed and the increase in the number of *E. coli* over time is presented in Fig. 2(c). The number of *E. coli* doubled constantly at intervals of 56 min, which is consistent with the generation time of *E. coli* at room temperature [49]. These results clearly validate the label-free imaging capability of ODT for studying the dynamics of individual bacteria.

**Fig. 2.**
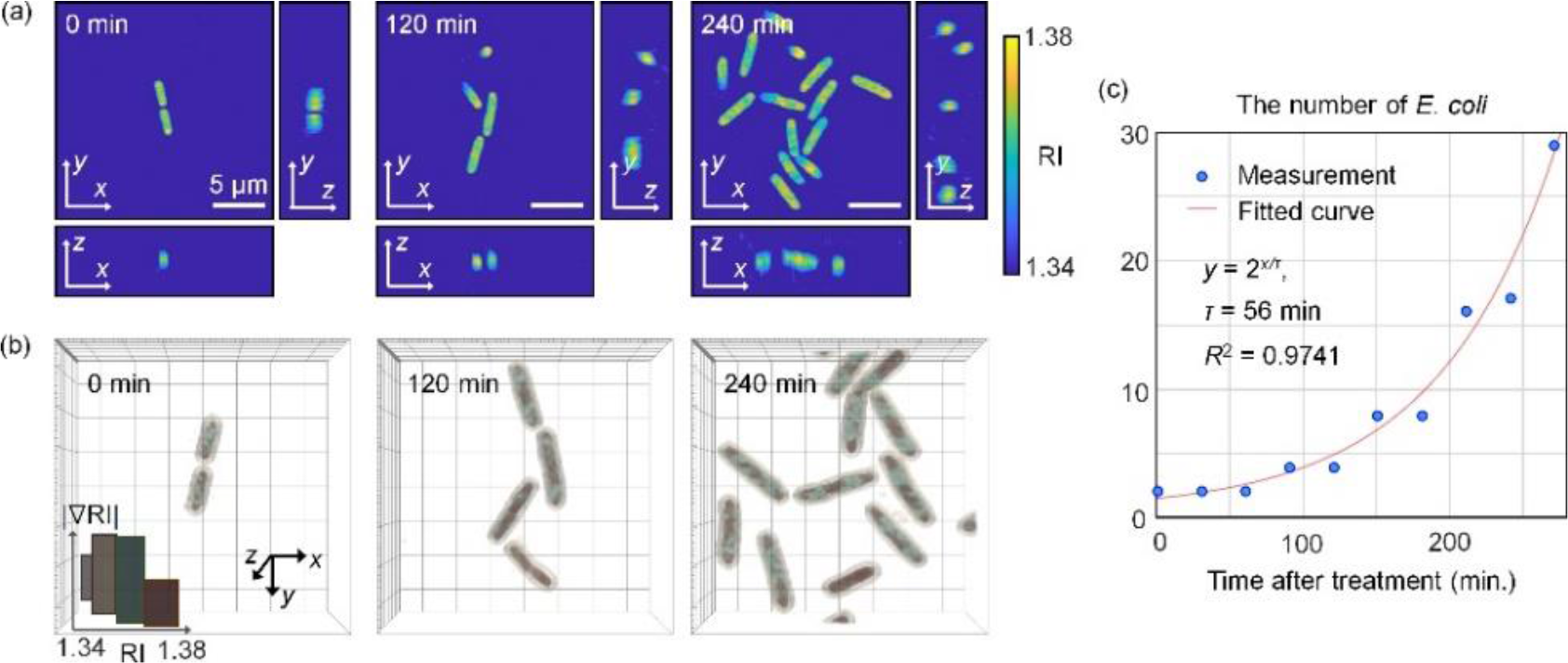
(a) Three-section views of the reconstructed RI distributions of *E. coli* without antibiotics over time (at 0, 120, and 240 minutes). (b) 3D rendered images of bacteria corresponding to (a). The color map shows various colors depending on the value of the RI. (c) The number of *E. coli* for the control group over time and the fitted curve. R-square = 0.9741.

### 3.2 Antibiotic responses of Escherichia coli and Bacillus subtilis

Next, we investigated the growth dynamics of individual bacteria under antibiotic treatments. As Gram-positive and Gram-negative models, *E. coli* and *B. subtilis* were selected, respectively. The bacteria were treated with ampicillin at various concentrations of 20, 100, and 300 μg/mL and imaged at time intervals of 30 minutes after drug treatment. As a normal control group, untreated bacterial samples were also measured at the same time intervals.

The results are shown in Fig. 3. In the absence of antibiotic treatment, both *E. coli* and *B. subtilis* exhibited active growth and division. When the bacteria were treated with antibiotics, each cell showed significant morphological alterations, and also presented a decrease in RI values, suggesting cell lysis. To effectively visualize these morphological alterations, the maximum RI projection of each 3D RI tomograms is presented for each group at various time points in Fig. 3.

**Fig. 3.**
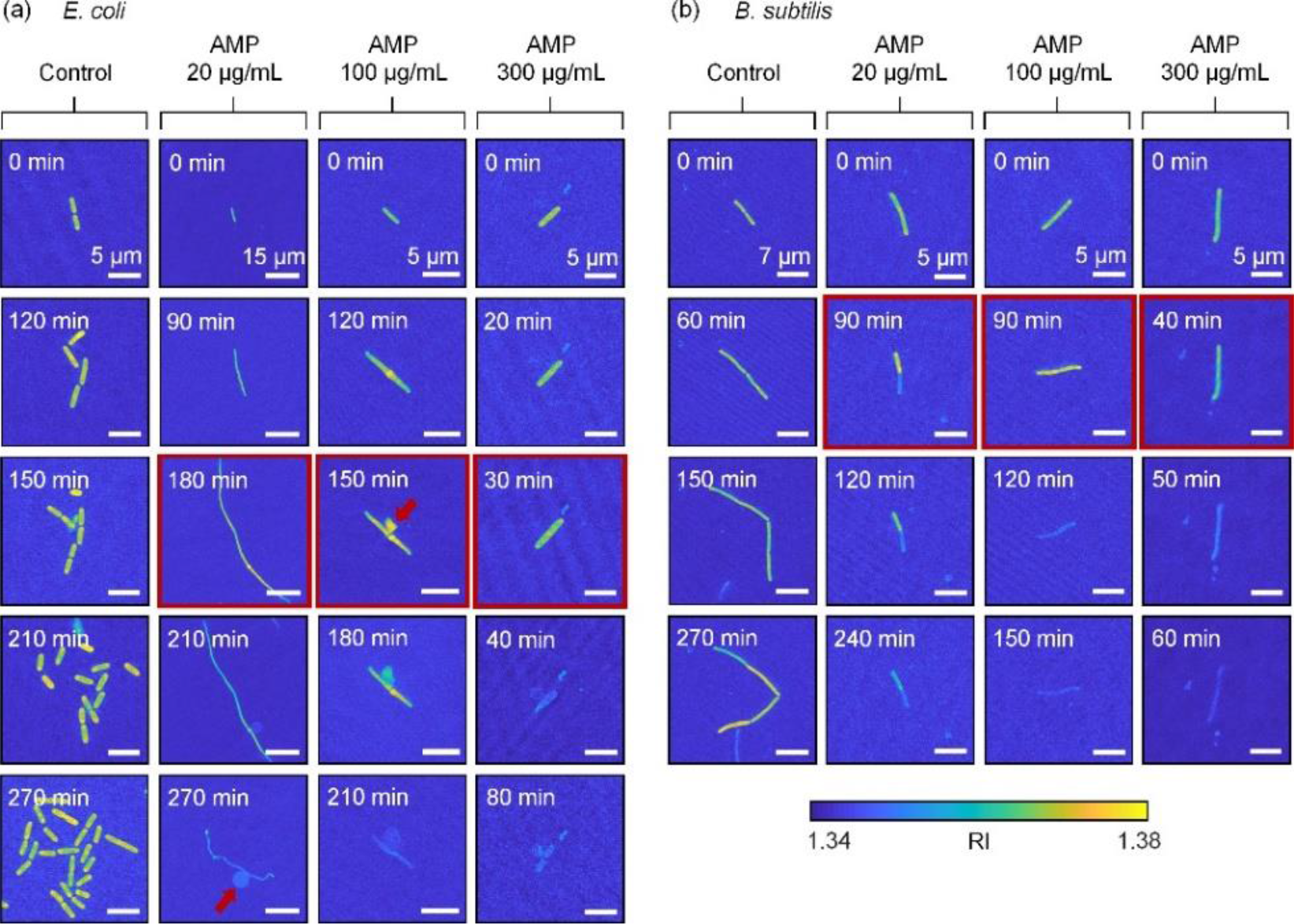
Maximum RI projection images with ampicillin over time. The red arrows indicate a bulge formed in *E. coli*. The entry with a red border is presumed to be near the point at which the bacteria die. (a) and (b) are *E. coli* and *B. subtilis*, respectively.

Figure 3(a) shows the morphological changes in *E. coli*, due to the ampicillin treatment. As the concentration of ampicillin increased, the bacteria underwent earlier cell lysis. From the reconstructed 3D RI tomograms, the point where the RI value drops suddenly could serve as the time point when the bacterial lysis initiated, because the decrease in the RI values of bacteria indicates the efflux of cell cytoplasm. When the concentration of the applied ampicillin was 20, 100, and 300 μg/mL, the bacterial lysis occurred right after 180, 150, and 30 minutes, respectively. In the control group, *E. coli* showed stable growth over time, whereas the bacteria treated with ampicillin did not divide.

In addition, the ampicillin-treated bacteria exhibited distinct elongation of cell shape; the cell length increased without cell division until the point when the RI value increased. Interestingly, the cells exposed to the ampicillin concentration of 20 μg/mL appeared like filaments, exhibiting a fiber-like cell shape. Notably, formation of bulges could be clearly detected in the measured tomograms [the red arrows in Fig. 3(a)]. Bulge formation was observed in groups treated with ampicillin concentrations of 20 or 100 μg/mL. However, a bulge was not formed at 300 μg/mL, which is a significantly high concentration. Furthermore, the rate of cell lysis increased noticeably as the concentration of ampicillin increased to 300 μg/mL.

Compared to the results with *E. coli*, *B. subtilis* exhibited significantly different responses to ampicillin [Fig. 3(b)]. First, the bacterial cell lysis time was earlier than that of *E. coli* treated with low ampicillin concentration. When the ampicillin concentration was 20, 100, and 300 μg/mL, the cell lysis points were conjectured to be right after 90, 90, and 40 minutes, respectively. At an even lower concentration, *B. subtilis* showed faster cell lysis than *E. coli*. In addition, *B. subtilis* exposed to the low concentration of ampicillin (20 μg/mL) did not exhibit morphological alteration into filament-like shapes as observed with *E. coli*. Notably, bulge formation was not observed in *B. subtilis*.

### 3.3 Quantitative analysis results for morphological and biochemical parameters

To investigate the morphological and biochemical characteristics of bacterial growth under antibiotic treatment, the following cellular parameters were retrieved and analyzed from the measured 3D RI tomograms: cell volume, cellular dry mass, cytoplasm concentration (See Methods). These cellular morphological and biochemical parameters were systematically addressed as a function of time and at various antibiotic concentrations [Fig. 4].

**Fig. 4.**
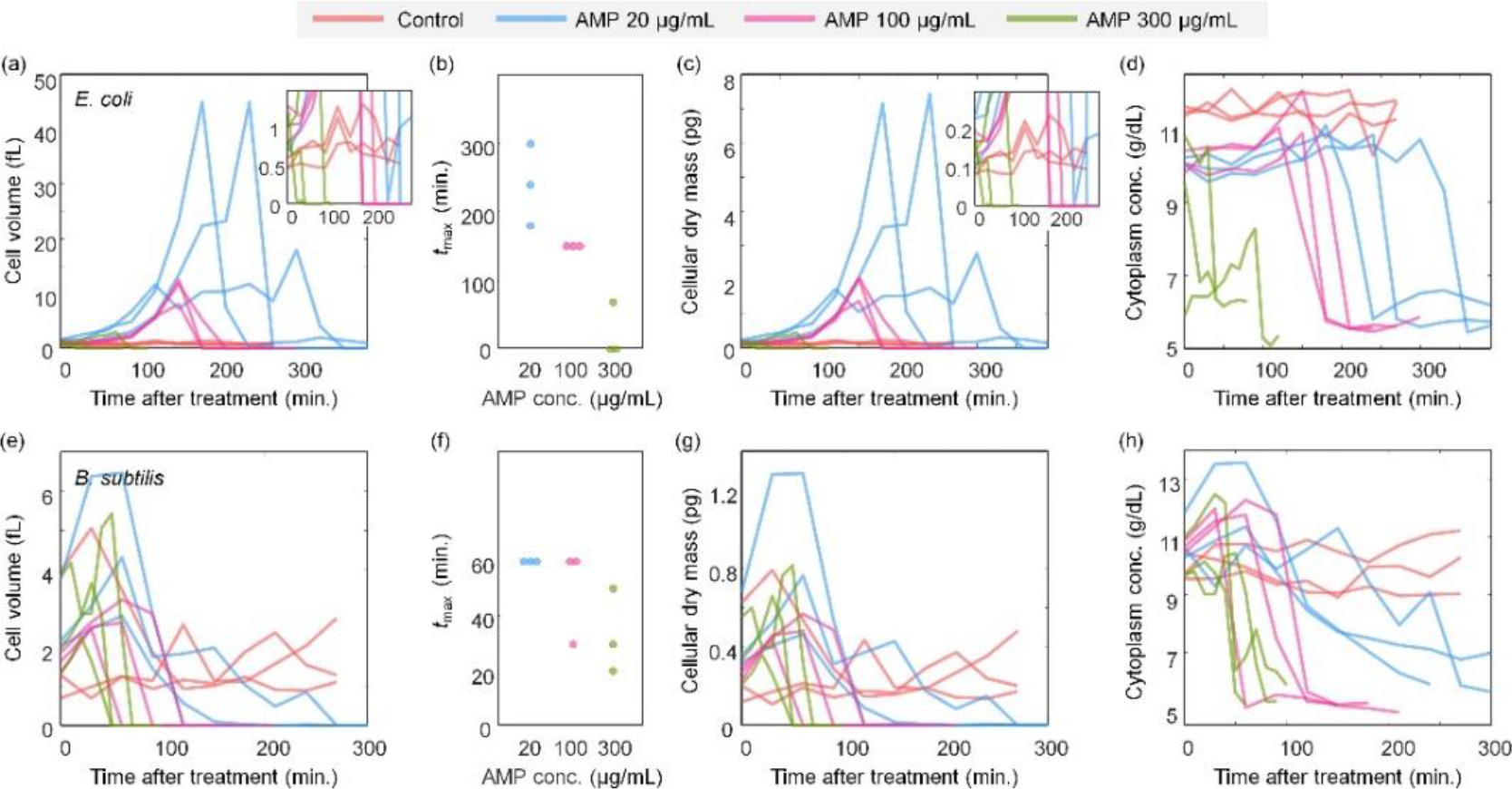
Cell volume, cellular dry mass, and cytoplasm concentration depending on the concentrations of ampicillin (0, 20, 100, and 300 μg/mL) and time at the maximum volume. (a–d) and (e–h) are *E. coli* and *B. subtilis*, respectively.

Figure 4(a) shows the cellular volume of *E. coli* over time for various concentrations of ampicillin. Note that this analysis was conducted at the single cell level, and each line in Fig. 4 indicates individual cells. For untreated *E. coli*, cells continued growth and division, and thus the volumes of each cell remained within a certain range (0.5–1.4 fL). Furthermore, the volumes of untreated *E. coli* oscillated over time, implying continuous growth and division.

Upon ampicillin treatment, *E. coli* did not exhibit cell division, as shown in Fig. 3. The volumes of treated *E. coli* showed monotonic increases, followed by sudden plummets [Fig. 3(a)]. These time points are consistent with the cell lysis observed in the 3D RI tomograms. Comparing the graphs for ampicillin concentrations of 20, 100, and 300 μg/mL, as the concentration of ampicillin increased, the bacteria showed earlier lysis, and the maximum volume of bacteria decreased. In addition, the maximum volume of *E. coli* with ampicillin treatments was significantly larger than that of the control group. These results
agree with our observation that when ampicillin is applied, cell division stops and abnormal elongation and bulge formation occurs. This abnormal growth occurred for a shorter time at high ampicillin concentration, indicating that *E. coli* burst to death faster at higher antibiotic concentrations [Fig. 4(b)].

The cellular dry mass of *E. coli* is shown in Fig. 4(c) (See Methods). The graphs for the dry mass are similar to those of cell volume [Fig. 4(a)], indicating that the cytoplasmic concentrations did not significantly change over time. This result can also be confirmed from the measured 3D RI tomograms [Fig. 4(d)]. The cytoplasm concentration increased slightly as the bacteria grew. After bacterial lysis, the internal contents escaped beyond the cell wall, and so, the cytoplasm concentration dropped suddenly. Comparison between the control and ampicillin-treated groups clearly indicates that the sudden drop in cytoplasm concentration was only notable in the latter case.

Compared to *E. coli*, *B. subtilis* did not show noticeable increases in either cell volume or dry mass upon ampicillin treatment [Figs. 4(e)-4(g)]. Rather, the bacteria rapidly reached a much smaller cell volume than the initial volume. The most discernible change was observed for the temporal graphs of cytoplasm concentration; a decrease in cytoplasm concentration caused by ampicillin was relatively well observed [Fig. 4(h)].

To clarify the different lysis behaviors between *E. coli* and *B. subtilis,* we reorganized and compared the species-dependent changes in quantitative parameters. Among the various parameters, we chose to compare cell volumes because they showed the most notable difference [Fig. 5]. At ampicillin concentrations of 20 and 100 μg/mL, the difference in cell volume between *E. coli* and *B. subtilis* was remarkable [Figs. 5(a) and 5(b)]. This is because abnormal growth was observed in *E. coli* treated with ampicillin, but no such growth was observed in *B. subtilis*. Moreover, at ampicillin concentrations of 20 and 100 μg/mL, the time to reach a minimal cell volume was shorter for *B. subtilis* than for *E. coli*. In other words, *B. subtilis* appeared to undergo lysis faster than *E. coli*. However, at the concentration of 300 μg/mL [Fig. 5(c)], this tendency was lost. These observations can be seen more clearly in Fig. 5(d), which represents the maximum volume of the bacteria at each concentration as a relative value. These data sufficiently represent how ODT can quantitatively address the dynamics of the antibiotic response of bacteria.

**Fig. 5.**
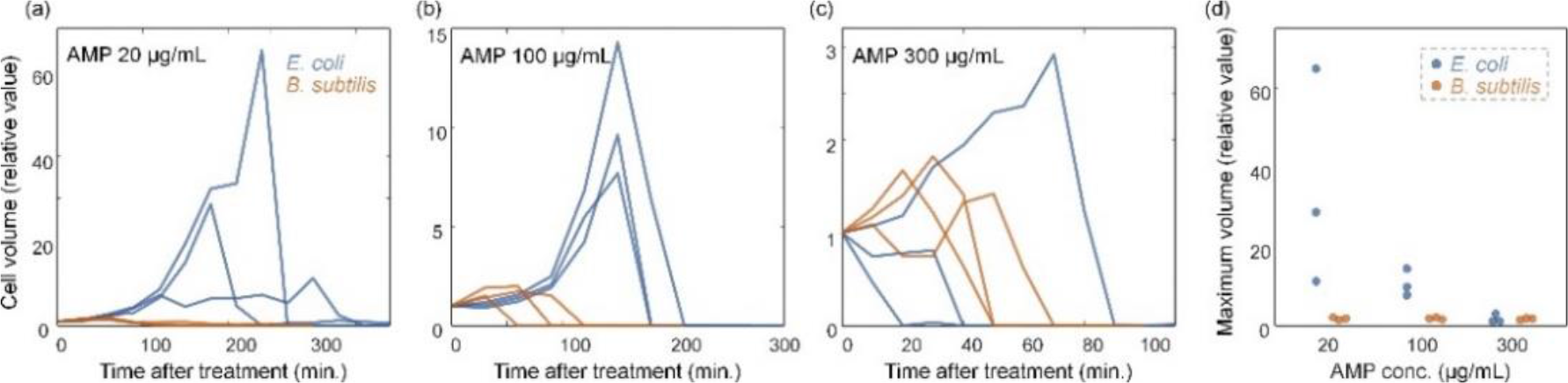
(a–c) The relative value of cell volume for *E. coli* and *B. subtilis* with the ampicillin concentration of (a) 20 μg/mL (b) 100 μg/mL (c) 300 μg/mL. (d) The relative value of the maximum volume for *E. coli* and *B. subtilis*.

## 4. Discussions and Summary

In summary, we present a method to systematically investigate the alterations in individual bacteria upon antibiotic treatment. Employing ODT, the 3D RI of antibiotic-treated bacteria was measured over time in a label-free and non-invasive manner. The 3D RI reconstruction of *E. coli* and *B. subtilis* bacteria provides the morphological and biochemical parameters related to their real-time alterations depending on the concentration of ampicillin, which were characterized by the inhibition of cell division, bulge formation, abnormal elongation, and cell lysis. Because the RI inside the bacteria is directly related to the cytoplasm concentration, dynamics of cytoplasmic concentrations and cellular dry mass were also analyzed quantitatively. Moreover, we quantitatively deduced the bacterial lysis time through a sudden change in cell volume or cellular dry mass. This can also be used effectively to study the cell cycle of bacteria [50].

Our measurements and observations with bacteria are consistent with those in the existing literature. Ampicillin from the β-lactam class inhibits cell wall synthesis, leading to various changes in bacterial cell shape and size, and finally resulting in bacterial cell death [51]. It is also intriguing to observe the changes in bacterial appearance depending on the ampicillin concentration. Spratt explains that cell division of *E. coli* is inhibited even at low concentrations of ampicillin, resulting in filamentation of *E. coli*; further, at higher concentrations, a bulge is formed in the middle of *E. coli*, and cell lysis occurs at significantly high ampicillin concentrations [13]. In our experiments, the corresponding concentrations can be considered as 20, 100, and 300 μg/mL, respectively. Comparing the cells with 20 μg/mL of ampicillin to the control group, cell division was inhibited, and abnormal growth, i.e. filamentation, was observed. As the concentration increased to 100 μg/mL, the filamentation phenomenon decreased, and the bulge became more pronounced. This is also consistent with the statement that only lysis was observed at the concentration of 300 μg/mL.

The different responses of *E. coli* and *B. subtilis* to ampicillin are also noteworthy for discussion. Ampicillin hinders transpeptidation by combining with penicillin-binding proteins on maturing peptidoglycan strands, and the resulting reduction in peptidoglycan synthesis causes cell lysis [13, 52]. In the case of Gram-positive bacteria including *B. subtilis*, the majority of the cell wall is composed of peptidoglycan; thus, the effect on ampicillin is more pronounced [52], resulting in faster cell lysis. This explains the contrast in the *E. coli* and *B. subtilis* at ampicillin concentrations of 20 and 100 μg/mL. In addition, there is a difference in the reactivity of Gram-positive and Gram-negative bacteria to ampicillin, but ampicillin eventually leads to cell death in both types of bacteria [53]. Our experiments confirmed that both *E. coli* and *B. subtilis* reached cell lysis in the presence of ampicillin. Indeed, our results were based on small datasets but were sufficient to epitomize the antibiotic response in a label-free, quantitative, and three-dimensional manner.

Previously, 2D QPI techniques have been applied for the study of individual bacteria [54], [55, 56]. However, 2D QPI only measures the optical phase delay maps, an integration of RI values along an optical path, and does not provide tomographic measurements of individual bacteria. As demonstrated in this work, ODT enables the systematic investigation of various volumetric information of individual bacteria. We emphasize that the 3D imaging technique exploited here is not limited to the objects targeted in this paper. The species of bacteria and the type of antibiotics can be expanded as needed. In addition, using this method, it is possible to conduct statistical analysis by increasing the number of microbes regardless of single cells or multiple cells. It is also possible to infer the physical functions and biological characteristics of bacteria from their interior structural features [57, 58]. Moreover, quantitative analysis for other morphological and biochemical parameters can be attempted. For instance, when a bulge is formed in *E. coli*, each volume or cytoplasm concentration can be quantified by separating the bulge and non-bulge portions. The lifetime of the bulge can also be measured. In this case, combination with the segmentation method will serve as the key to analysis [24]. We believe that the methods presented in this paper will provide microbiologists with broader insights into the various responses of bacteria to antibiotics.

## Acknowledgments

The authors thank Dr. Weisun Park for helpful discussions.

## References

1. Kohanski, M.A., D.J. Dwyer, and J.J. Collins, How antibiotics kill bacteria: from targets to networks. Nature Reviews Microbiology, 2010. 8(6): p. 423.

2. Pambos, O.J. and A.N. Kapanidis, Tracking antibiotic mechanisms. Nature Reviews Microbiology, 2019. 17(4): p. 201–201.

3. Aminov, R.I., A brief history of the antibiotic era: lessons learned and challenges for the future. Frontiers in microbiology, 2010. 1: p. 134–134.

4. Neu, H.C., The crisis in antibiotic resistance. Science, 1992. 257(5073): p. 1064–1073.

5. Conly, J. and B. Johnston, Where are all the new antibiotics? The new antibiotic paradox. Canadian Journal of Infectious Diseases and Medical Microbiology, 2005. 16(3): p. 159–160.

6. Veses Garcia, M., et al., Rapid Phenotypic Antibiotic Susceptibility Testing of Uropathogens Using Optical Signal Analysis on the Nanowell Slide. Frontiers in microbiology, 2018. 9: p. 1530.

7. Malmberg, C., et al., A novel microfluidic assay for rapid phenotypic antibiotic susceptibility testing of bacteria detected in clinical blood cultures. PloS one, 2016. 11(12): p. e0167356.

8. Choi, J., et al., A rapid antimicrobial susceptibility test based on single-cell morphological analysis. Science translational medicine, 2014. 6(267): p. 267ra174–267ra174.

9. Fredborg, M., et al., Rapid antimicrobial susceptibility testing of clinical isolates by digital time-lapse microscopy. European Journal of Clinical Microbiology & Infectious Diseases, 2015. 34(12): p. 2385–2394.

10. Fredborg, M., et al., Real-time optical antimicrobial susceptibility testing. Journal of clinical microbiology, 2013. 51(7): p. 2047–2053.

11. Qi, C., C.W. Stratton, and X. Zheng, Phenotypic testing of bacterial antimicrobial susceptibility, in Advanced Techniques in Diagnostic Microbiology. 2006, Springer. p. 63–83.

12. Bochner, B.R., L. Giovannetti, and C. Viti, Important discoveries from analysing bacterial phenotypes. Molecular Microbiology, 2008. 70(2): p. 274–280.

13. Spratt, B.G., Distinct penicillin binding proteins involved in the division, elongation, and shape of Escherichia coli K12. Proceedings of the National Academy of Sciences, 1975. 72(8): p. 2999–3003.

14. Yao, Z., D. Kahne, and R. Kishony, Distinct single-cell morphological dynamics under beta-lactam antibiotics. Molecular cell, 2012. 48(5): p. 705–712.

15. Lewis, C.L., C.C. Craig, and A.G. Senecal, Mass and density measurements of live and dead gram-negative and gram-positive bacterial populations. Appl. Environ. Microbiol., 2014. 80(12): p. 3622–3631.

16. Delgado, F.F., et al., Intracellular water exchange for measuring the dry mass, water mass and changes in chemical composition of living cells. PloS one, 2013. 8(7): p. e67590.

17. Jesacher, A., M. Ritsch-Marte, and R. Piestun, Three-dimensional information from two-dimensional scans: a scanning microscope with postacquisition refocusing capability. Optica, 2015. 2(3): p. 210–213.

18. Jensen, E.C., Use of fluorescent probes: their effect on cell biology and limitations. The Anatomical Record: Advances in Integrative Anatomy and Evolutionary Biology, 2012. 295(12): p. 2031–2036.

19. Murtey, M.D. and P. Ramasamy, Sample preparations for scanning electron microscopy–life sciences, in Modern electron microscopy in physical and life sciences. 2016, IntechOpen.

20. Park, Y., C. Depeursinge, and G. Popescu, Quantitative phase imaging in biomedicine. Nature Photonics, 2018. 12(10): p. 578–589.

21. El Kirat, K., et al., Sample preparation procedures for biological atomic force microscopy. Journal of microscopy, 2005. 218(3): p. 199–207.

22. Koo, S.-e., et al., Reconstructed Three-Dimensional Images and Parameters of Individual Erythrocytes Using Optical Diffraction Tomography Microscopy. Annals of laboratory medicine, 2019. 39(2): p. 223–226.

23. Cho, C., et al., Study of Optical Configurations for Multiple Enhancement of Microalgal Biomass Production. Scientific reports, 2019. 9(1): p. 1723.

24. Lee, M., et al., Deep-learning based three-dimensional label-free tracking and analysis of immunological synapses of chimeric antigen receptor T cells. BioRxiv, 2019: p. 539858.

25. Tougan, T., et al., Molecular camouflage of Plasmodium falciparum merozoites by binding of host vitronectin to P47 fragment of SERA5. Scientific reports, 2018. 8(1): p. 5052.

26. Park, H., et al., Characterizations of individual mouse red blood cells parasitized by Babesia microti using 3-D holographic microscopy. Scientific reports, 2015. 5: p. 10827.

27. Park, C., et al., Three-dimensional refractive-index distributions of individual angiosperm pollen grains. Current Optics and Photonics, 2018. 2(5): p. 460–467.

28. Oh, S., et al., In situ measurement of absolute concentrations by Normalized Raman Imaging. BioRxiv, 2019: p. 629543.

29. Yang, S.A., et al., Measurements of morphological and biophysical alterations in individual neuron cells associated with early neurotoxic effects in Parkinson’s disease. Cytometry part A, 2017. 91(5): p. 510–518.

30. Kwon, S., et al., Mitochondria-targeting indolizino [3, 2-c] quinolines as novel class of photosensitizers for photodynamic anticancer activity. European journal of medicinal chemistry, 2018. 148: p. 116–127.

31. Liu, Y.P., Refractive Index Distribution of Single Cell and Bacterium Usingan Optical Diffraction Tomography System. 2016.

32. Lauer, V., New approach to optical diffraction tomography yielding a vector equation of diffraction tomography and a novel tomographic microscope. Journal of Microscopy, 2002. 205(2): p. 165–176.

33. Shin, S., et al., Active illumination using a digital micromirror device for quantitative phase imaging. Optics Letters, 2015. 40(22): p. 5407–5410.

34. Wolf, E., Three-dimensional structure determination of semi-transparent objects from holographic data. Optics Communications, 1969. 1(4): p. 153–156.

35. Kim, K., et al., Optical diffraction tomography techniques for the study of cell pathophysiology. Journal of Biomedical Photonics & Engineering, 2016. 2(2).

36. Takeda, M., H. Ina, and S. Kobayashi, Fourier-transform method of fringe-pattern analysis for computer-based topography and interferometry. JosA, 1982. 72(1): p. 156–160.

37. Debnath, S.K. and Y. Park, Real-time quantitative phase imaging with a spatial phase-shifting algorithm. Optics letters, 2011. 36(23): p. 4677–4679.

38. Sung, Y., et al., Optical diffraction tomography for high resolution live cell imaging. Optics express, 2009. 17(1): p. 266–277.

39. Kim, K., et al., High-resolution three-dimensional imaging of red blood cells parasitized by Plasmodium falciparum and in situ hemozoin crystals using optical diffraction tomography. Journal of biomedical optics, 2013. 19(1): p. 011005.

40. Devaney, A., Inverse-scattering theory within the Rytov approximation. Optics letters, 1981. 6(8): p. 374–376.

41. Devaney, A.J., Mathematical foundations of imaging, tomography and wavefield inversion. 2012: Cambridge University Press.

42. Park, C., S. Shin, and Y. Park, Generalized quantification of three-dimensional resolution in optical diffraction tomography using the projection of maximal spatial bandwidths. JOSA A, 2018. 35(11): p. 1891–1898.

43. Lim, J., et al., Comparative study of iterative reconstruction algorithms for missing cone problems in optical diffraction tomography. Optics express, 2015. 23(13): p. 16933–16948.

44. Gerchberg, R., Super-resolution through error energy reduction. Optica Acta: International Journal of Optics, 1974. 21(9): p. 709–720.

45. Papoulis, A., A new algorithm in spectral analysis and band-limited extrapolation. IEEE Transactions on Circuits and systems, 1975. 22(9): p. 735–742.

46. Barer, R. and S. Joseph, Refractometry of living cells: Part I. Basic principles. Journal of Cell Science, 1954. 3(32): p. 399–423.

47. Popescu, G., et al., Optical imaging of cell mass and growth dynamics. American Journal of Physiology-Cell Physiology, 2008. 295(2): p. C538–C544.

48. Jung, J., et al., Label-free non-invasive quantitative measurement of lipid contents in individual microalgal cells using refractive index tomography. Scientific reports, 2018. 8.

49. Marr, A.G., Growth rate of Escherichia coli. Microbiology and Molecular Biology Reviews, 1991. 55(2): p. 316–333.

50. Van Iterson, W., Some features of a remarkable organelle in Bacillus subtilis. The Journal of Cell Biology, 1961. 9(1): p. 183–192.

51. Donachie, W.D., The cell cycle of Escherichia coli. Annual review of microbiology, 1993. 47(1): p. 199–230.

52. Efstratiou, E., et al., Antimicrobial activity of Calendula officinalis petal extracts against fungi, as well as Gram-negative and Gram-positive clinical pathogens. Complementary Therapies in Clinical Practice, 2012. 18(3): p. 173–176.

53. Percival, A., W. Brumfitt, and J. De Louvois, The role of penicillinase in determining natural and acquired resistance of gram-negative bacteria to penicillins. Microbiology, 1963. 32(1): p. 77–89.

54. Kemper, B., et al., Towards 3D modelling and imaging of infection scenarios at the single cell level using holographic optical tweezers and digital holographic microscopy. Journal of biophotonics, 2013. 6(3): p. 260–266.

55. Jo, Y., et al., Angle-resolved light scattering of individual rod-shaped bacteria based on Fourier transform light scattering. Scientific reports, 2014. 4: p. 5090.

56. Alanazi, H., et al., Robust microbial cell segmentation by optical-phase thresholding with minimal processing requirements. Cytometry Part A, 2017. 91(5): p. 443–449.

57. Tanaka, S., M.R. Sawaya, and T.O. Yeates, Structure and mechanisms of a protein-based organelle in Escherichia coli. Science, 2010. 327(5961): p. 81–84.

58. Parsons, J.B., et al., Biochemical and structural insights into bacterial organelle form and biogenesis. Journal of biological chemistry, 2008. 283(21): p. 14366–14375.

